# From local motifs to global dynamical stability in the mouse brain connectome

**DOI:** 10.64898/2026.02.27.708665

**Authors:** Yuze Liu, Yina Wei, Hanchuan Peng

**Affiliations:** New Cornerstone Science Laboratory, SEU-ALLEN Joint Center, Institute for Brain and Intelligence, Southeast University, Nanjing, China; New Cornerstone Science Laboratory, Institute for Brain and Intelligence, Fudan University, Shanghai, China; Shanghai Academy of Natural Sciences (SANS), Fudan University, Shanghai, China

## Abstract

Brain connectomics provides insights into the architectural organization of neural circuits across the brain. But it remains unclear how neurons and their connections give rise to brain-wide activity. Here, using a sparse detailed mouse brain connection map called “bouton-net,” we studied how specific neuron patterns (motifs) and groups (modules) affect information processing. We discovered that complex, “two-way” connections are much more common than expected. These patterns appear across all brain regions, suggesting they are a universal rule for brain wiring. We also found that different brain modules do not communicate randomly; instead, they talk through a small group of highly connected neurons. By testing these organization patterns with computer models, we found that the brain may not be wired for the best memory capacity or nonlinear processing capability, but to ensure that capability remains stable. This structure provides the foundation for a reliable, “crash-proof” biological network.

## Introduction

Understanding how neural connectivity constrains brain organization and dynamics is a central challenge in neuroscience. The brain can be conceptualized as a complex network of neurons and synaptic connections, a perspective formalized by network neuroscience, which applies graph-theoretical approaches to relate connectivity with function [1]. Across species and scales, biological connectomes exhibit some topological features, including modular organization, hub nodes, and recurrent connectivity, which can be used to distinguish biological networks from random networks and are thought to support efficient and robust information processing [2,3,4].

Network motifs, defined as small subgraph patterns that occur more frequently than expected by chance, provide a framework for characterizing higher-order connectivity structure [5]. Previous works demonstrated that specific motif classes, such as feedforward loops and reciprocal connections, are selectively enriched in biological networks, suggesting functional constraints on network architecture [5,6,7,8]. In local circuit neural networks of rat, directed motifs, particularly bidirectional connections and three-neuron patterns, have been associated with synaptic connection strength distribution, which implied the correlation between structural and functional connectomes [9]. However, how such motifs distribute across large-scale modular organization, especially at single-neuron resolution, remains unclear.

At the mesoscale, brain networks are organized into modules that support functional specialization while remaining integrated through inter-modular connections [10]. Hub nodes and rich-club structures play a critical role in mediating cross-module communication and shaping global dynamics [11,12]. Although modularity and motif structure have each been linked to network function, their interplay, particularly how motif organization aligns with modular directionality and neuron-level connectivity, has not been systematically examined in whole-brain biological networks.

Recent advances in electron microscopy (EM) connectomics have revealed non-random motif structure and recurrent connectivity in local circuits, including insect connectomes and mammalian cortical volumes [13,14,15,16]. Large-scale reconstruction efforts, such as those from the MICrONS program and the Blue Brain Project, demonstrate that biologically constrained microcircuits exhibit rich higher-order connectivity patterns that shape emergent activity [17,18,19]. However, these datasets are spatially limited, leaving unresolved whether motif organization observed locally is conserved across the entire brain and how it constrains global network dynamics.

Another open question concerns the functional consequences of motif and modular organization. While connectomic data increasingly resolve synapse-level wiring, deriving nonlinear dynamical properties directly from network topology remains challenging [20]. By driving fixed, untrained recurrent networks with time-varying inputs, reservoir computing elicits rich transient dynamics, thereby providing a computational framework to bridge this gap [21,22], enabling systematic evaluation of how recurrent connectivity patterns influence memory, stability, and temporal processing [23,24].

Here, we address these challenges using bouton-net, a sparse brain-wide, single-neuron level mouse connectome [25]. We show that complex, bidirectionally connected triadic motifs are strongly enriched relative to random networks and are consistently preserved across modules composed of distinct brain regions. We further demonstrate that modules exhibit distinct directional preferences in inter-modular connectivity, and that inter-modular projections contribute disproportionately to diverse motif classes, often mediated by a limited subset of high-degree neurons. Using a reservoir computing framework, we show that although the biological network does not maximize task performance, it exhibits enhanced dynamical stability compared with random networks. Perturbation of bidirectional connections reduces motif occurrence, directly linking single-neuron motif organization to stable network dynamics.

## Results

### Structural motif reveals the module complexity of biological brain networks

We used a mouse whole-brain connectome dataset, bouton-net, consisting of 1,877 neurons with complete reconstructions of their morphology, registered in the Allen CCFv3 framework. The dataset spans 90 brain regions, with major contributions from thalamus (VPM, ventral posteromedial; 389 neurons), striatum (CP, caudoputamen; 324 neurons), and sensorimotor cortex (SSp-m, mouth primary somatosensory; 87 neurons), alongside representatives from visual, auditory, and hippocampal areas. Neuronal connections were computed based on a nuanced Peters’ rule that accounts for axonal bouton location as a proxy of synaptic connectivity beyond spatial proximity. Based on a graph segmentation algorithm, the bouton-net was partitioned into 12 modules (B1–B12), comprising 1,210 neurons in total, with each module containing more than 30 neurons. These modules exhibit significantly higher intra-modular connectivity density compared to inter-modular connectivity density [25], with neurons in each module relatively spatially clustered in close proximity (**Fig. 1A**).

**Figure 1.**
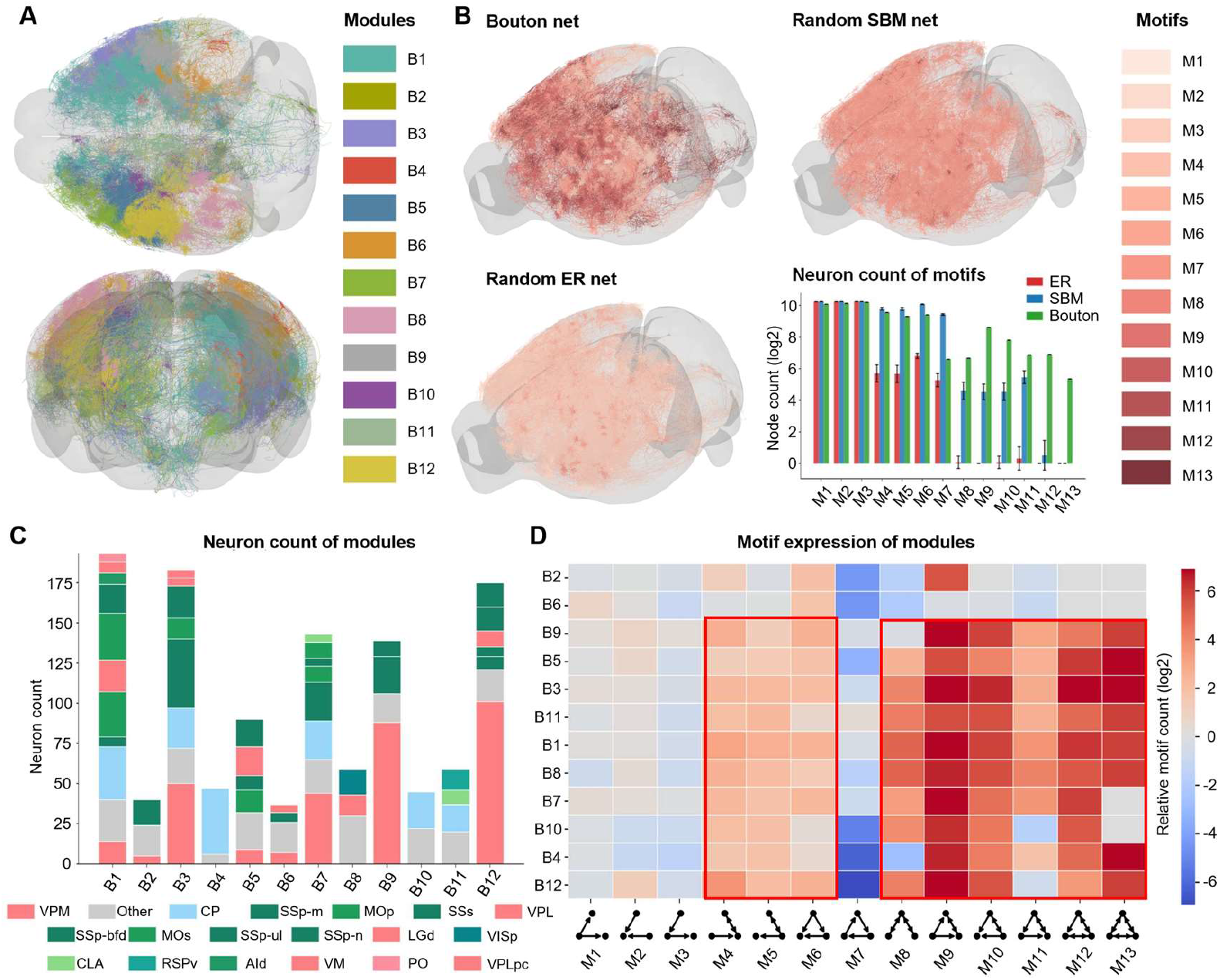
The occurrence pattern of triad motifs in whole bouton-net and each module of mouse brain. **A**, Neurons spatial distribution of 12 modules in bouton-net (1210 neurons in total). Colors indicate different modules. **B**, Comparison of motifs’ spatial distribution of neurons and the number of neurons which participant in motifs between bouton-net, ER (Erdős–Rényi) random net and SBM (stochastic block model) random net. Color bar, the level of motifs in triad census. The motif identity of each neuron is defined as its involvement of the highest motif. The error bars represent the mean neuron numbers±standard deviation. **C**, Neuron numbers of each module in mouse brain. Colors indicate different brain regions according to Allen Brain Atlas. **D**, Occurrence ratio of 13 motifs in different modules of bouton-net compared to ER random networks. The red boxes highlight the motifs which have over-representation. Color bar, the number of motifs counts ratio (log2 transformed).

To assess network topology, we conducted a triad census using 13 non-redundant motifs, each containing three nodes with no isolated elements (M1–M13). This triad census was applied to bouton-net as well as several randomized networks which were driven by specific hyperparameters to maintain the same network scale (**Methods**), including an Erdős–Rényi (ER) random network [26], Scale-Free (SF) random network [27] and a random Stochastic Block Model (SBM) network [28] which mimics the biological module structures. To visualize structural patterns, nodes were colored according to their involvement of the highest motif level (M1-M13), allowing for a direct comparison of participation across network types (**Fig. 1B**). Compared with the bouton-net (**Fig. 1B, top left**), while the ER network (**Fig. 1B, bottom left)** was dominated by elementary triadic interactions (M1–M7) **(Fig. 1B, bottom right, red**). Although the SF network has significant positive correlation with bouton-net in terms of the degree distribution [25], it has similar prevalences of elementary structural motifs to those found in the ER network. Especially SF has a relatively large number of feed-forward loops (M6) (**Supplementary Fig 1, red**). The biologically informed SBM (**Fig. 1B, top right**) more closely approximated the bouton-net (**Fig. 1B, bottom right, green**) by generating a motif distribution characteristic of complex module systems **(Fig. 1B, bottom right, blue)**. Notably, the SBM preserved the representation of basic motifs while significantly elevating the prevalence of complex **(Fig. 1B, bottom right, blue)**, bidirectional-connection structures (M8–M12). However, even at a constant network size, the bouton-net consistently exhibited a substantially higher node count within these complex motifs than SBM, highlighting a remaining gap in structural density.

These observations underscore that triad census motifs can effectively distinguish between artificial networks and real biological networks, revealing distinct topologies across various types of constructed networks.

### Consistent motif occurrence across modules in bouton-net

To further investigate whether motif occurrence patterns are conserved across different modules within biological networks, we analyzed the 12 bouton-net modules based on their brain region composition, and network size (**Fig. 1C**). Although all modules exhibited high connectivity density, the number of nodes per module varied considerably, likely due to the sparse nature of the bouton-net. Additionally, while the modules contained neurons from diverse brain regions, the upstream and downstream brain area analysis of each module suggested functional specialization.

We conducted triad motif censuses for each module and compared the frequency of motif occurrences to a randomized ER network of equivalent size, calculating the motif occurrence ratio (**Fig. 1D, Methods**). Most modules exhibited consistent over-representation of motifs M4–M6 and M8–M13, relative to the random ER network. These motifs were characterized by bidirectional connections, indicating significant non-random interactions between neurons. However, modules B2 and B6 displayed under-representation of these motifs, suggesting a possible influence of their smaller module sizes and the inherent sparsity of the connectome data, which could introduce sampling irregularities.

Overall, triad motif census reveals notable differences between bouton-net and random ER networks. Furthermore, the motif occurrence patterns are consistent across modules of varying network sizes and node compositions of brain regions. The bidirectional connections observed in overexpressed motifs suggest that neuronal connectivity in biological brain networks is far from random, potentially reflecting underlying specific advantages of function.

### Directional preferences of information transmission across network modules

We analyzed the directional connectivity preferences of the modules in bouton-net by calculating the total number of outward and inward connections between each module’s nodes and those of other modules. These were defined as outward-modular connections and inward-modular connections, respectively. We averaged the number of connections per node within each module to quantify the preferential directionality of information transmission or reception across different modules (**Fig. 2A**). To mitigate potential sampling biases due to the sparsity of bouton-net and variations in module size, we randomly sampled 90% nodes within each module. After repeated sampling, we found that different modules exhibited distinct directional preferences in their connections. For example, the B5 module showed a significant preference for outward-modular connections, with fewer inward-modular connections, while the B10 module displayed the opposite pattern. The B8 module, an extreme case, primarily consisted of neurons from the visual cortex (VISp, Primary Visual Cortex; 35 neurons) and its subregions, which are typically low in number in bouton-net. As a high-level information processing brain area, this module exhibited reduced inter-modular connections, likely due to its specialized function within the brain’s circuitry.

**Figure 2.**
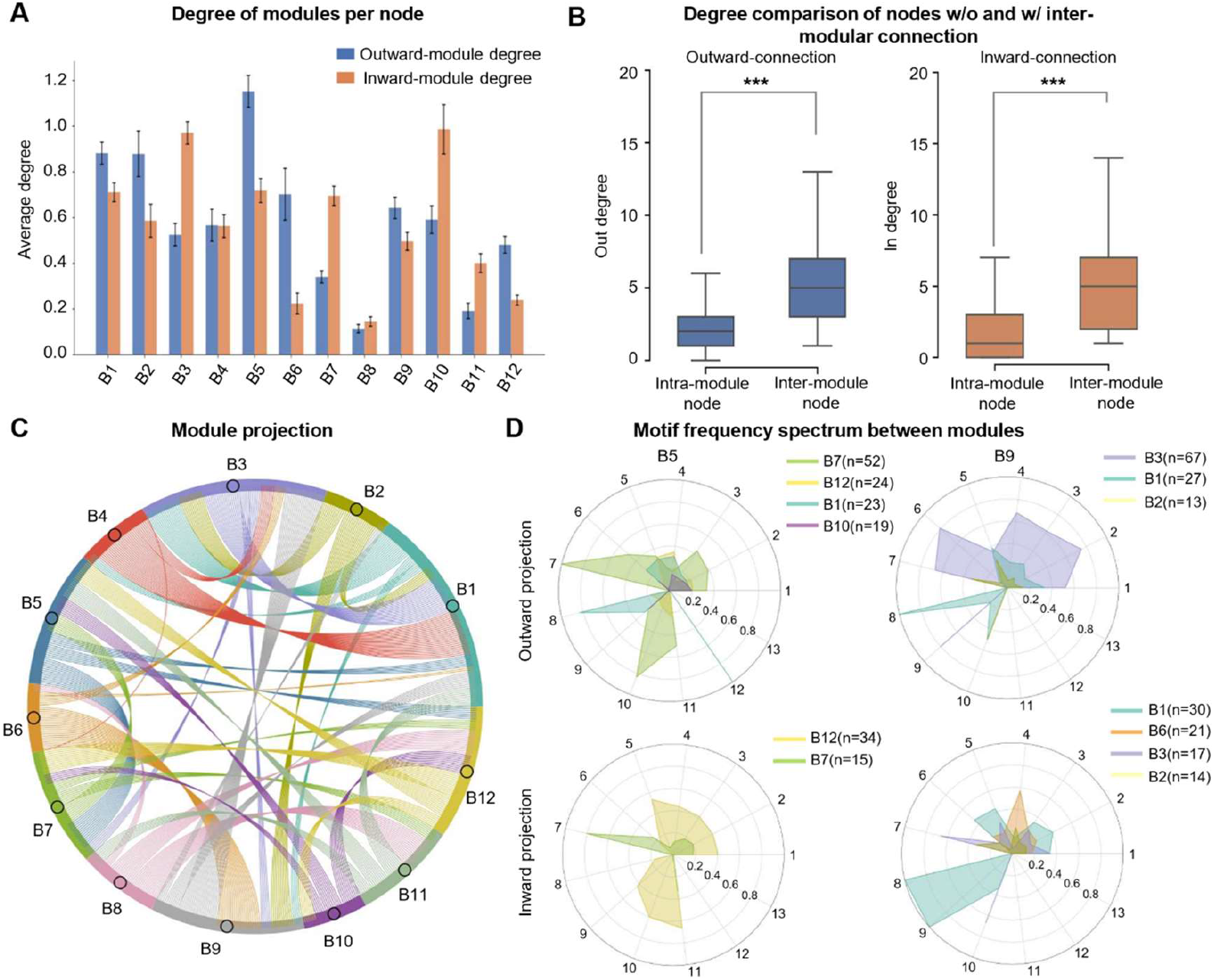
Directional projection preferences and motif contributions of module in bouton-net. **A**, Outward-modular/inward-modular degree of modules in bouton-net, which describes the number of edges that each module projects outward and receives from the outside. Colors indicate the types of degree. The error bars represent the mean degrees±standard deviation. **B**, Distribution of out-degree and in-degree for nodes with inter-modular connections (outward-module or inward-module) versus nodes with intra-modular connections. Colors indicate out-degree (blue) and in-degree (orange). Statistical significance was assessed using two-tailed Mann–Whitney U tests, *** represents p < 0.001. **C**, Outward projection pattern between modules in bouton-net. Colors indicate different brain modules. **D**, Outward and inward motif frequency spectrum of module B5 and B9, which demonstrate the proportion of motif composition of inter-modular connections between different modules. Colors indicate brain modules.

These findings suggest that, within biological networks like bouton-net, different modules exhibit varying directional preferences in their inter-modular connections. These preferences may indicate the roles that each module plays in functional circuits and their specific contribution to information flow during complex brain functions.

### Hub nodes preferentially support inter-modular information transmission

To explore whether the distribution of intra-modular and inter-modular connections among nodes is uniform, we distinguished nodes with only intra-modular connections from those with inter-modular connections, further subdividing the inter-modular connections into outward and inward categories to represent the directionality of inter-modular communication (**Fig. 2B**). We then statistically tested the distribution of out-degrees and in-degrees for nodes contributing to outward-modular and inward-modular connections respectively. We observed that nodes with outward-modular connections had significantly higher out-degrees than the rest (**Fig. 2B, blue**, the Mann-Whitney U test, two-sided, p = 2.77e-78), and nodes with inward-modular connections had higher in-degrees compared to others (**Fig. 2B, orange**, the Mann-Whitney U test, two-sided, p = 2.94e-82). This indicates that the internal and external connections within a modular are not uniformly distributed; rather, they are significantly biased toward high-degree hub nodes.

Thus, in bouton-net, nodes contributing to outward-modular or inward-modular connections tend to have larger degrees, suggesting that hub nodes not only serve as central connectors within their modules but also play pivotal roles in inter-modular information transmission.

### Inter-modular connectivity preferences and motif diversity in bouton-net

Given the directional property of neuronal connections, we further analyzed the projection preferences between modules by calculating the proportion of projections between modules and examining whether these projections contribute to specific motifs. We found that the projections between modules are not uniformly distributed but exhibit directional preferences (**Fig. 2C**). Despite the diverse brain region composition within each module, these projections reflect functional circuit-driven directionality, suggesting that different modules represent distinct brain functions and communicate through specific connection patterns.

We focused on the B5 and B9 modules, which are involved in sensory and motor functions [25], and analyzed their outward and inward connections in terms of their contribution to various motifs, considering network size and motif occurrence consistency (**Fig. 2D**). After excluding cases where small edge numbers (edge < 10) led to the formation of specific motifs, we found that for outward-modular connections, the B5 module and its functionally related B12 module showed relatively low occurrence of most motifs. In contrast, the more complex motifs (M10, M11) were predominantly formed by projections directed toward B7, likely due to the directional nature of the functional brain circuits. For inward-modular connections, B5 showed a predominant connection to B12, with these connections forming complex motifs such as M9, M10, and M11, in addition to simpler motifs (M1-M5). On the other hand, B9 exhibited outward connections with B3, forming simpler motifs, and both outward- and inward-modular connections with B1 contributed to complex motifs, particularly those involving M9, M10, and M11. Given that both B1 and B3 contain the CP (cerebellar) brain region, these connections may be related to motor cortex and caudate-putamen nuclei signaling in motor control and feedback mechanisms.

In conclusion, we find that projections between modules exhibit preferences consistent with functional specialization of the circuits, and these projections contribute to a diverse range of motifs, highlighting the complexity of information transfer between different brain regions.

### Stability of bouton-net during functional tasks

We adopted the reservoir computing framework and used Memory capacity and nonlinear autoregressive moving-average (NARMA-n) [21] tasks (n = 5, 10, 20) to assess the information storage ability and nonlinear processing capability of networks. The bouton-net, size-matched ER random networks, and several topology-perturbed (perturb bidirectional connections) networks based on bouton-net were assigned with random connection weights, were used as reservoirs (**Fig. 3A; Methods**). The spectral radius, defined as the largest absolute eigenvalue of the network weight matrix, serves as a critical control parameter in reservoir computing. By scaling spectral radius of network, it can regulate the reservoir’s dynamical regime to stable, contractive dynamics or chaotic, fading-memory dynamics. We systematically varied the spectral radius within the range of 0–2 to assess its impact on task performance across different dynamical regimes.

**Figure 3.**
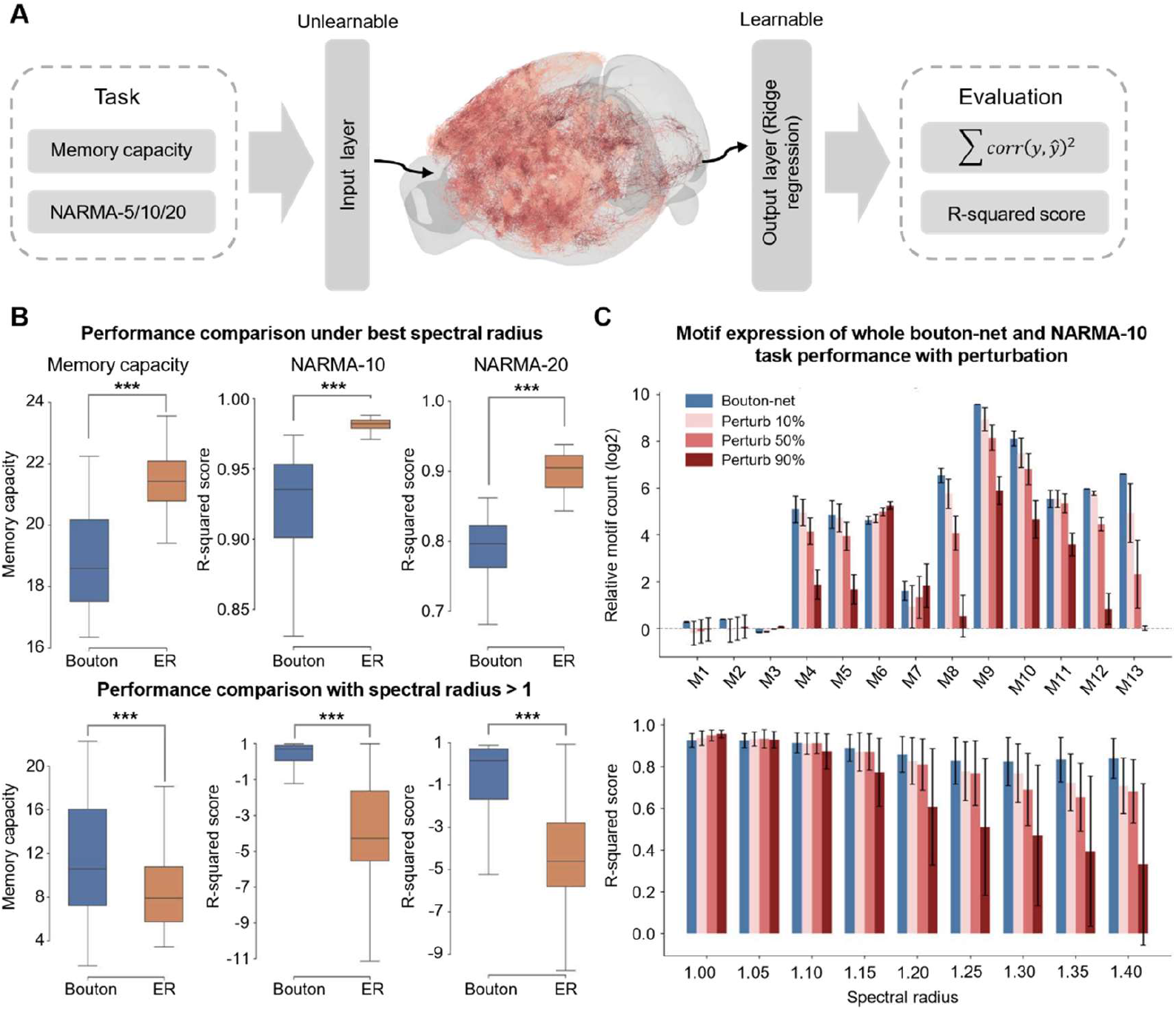
Stability-performance trade-off and motif-dependent stability in bouton-net. **A**, Schematic of the reservoir computing framework. Bouton-net, size-matched ER random networks, and topology-perturbed variants of bouton-net were used and evaluated on Memory capacity and NARMA-n (n=5, 10, 20) tasks. **B**, Performance comparison of tasks between bouton net and ER random net with increasing spectral radius. Top: distribution comparison of performance at the spectral radius with best task performance. Bottom: distribution comparison of performance with spectral radius above 1. Statistical significance was assessed using two-sided Mann–Whitney U tests, *** represents p < 0.001. Colors indicate the types of networks. **C**, Relationship between motif occurrence and network stability. Top: motif counts ratio (log2 transformed) of bouton-net and its perturbed variants relative to ER random networks, across different bidirectional connection perturbation proportions. The error bars represent the mean ratios±standard deviation. Bottom: NARMA-10 task performance of networks under chaotic regime. Colors indicate the original bouton-net and networks variants with increasing disrupting proportions. The error bars represent the mean R-squared scores±standard deviation.

We compared bouton-net (**Fig. 3B, blue**) with ER random networks (**Fig. 3B, orange**) under identical task conditions. In the Memory capacity task, bouton-net exhibited significantly lower peak performance than the ER random network at the optimal spectral radius (**Fig. 3B, top left;** the Mann-Whitney U test, two-sided, p = 3.42e-12). This result is consistent with previous findings about reciprocity, suggesting that the elevated reciprocity observed in biological networks may reduce effective memory capacity [29]. Since reciprocity reflects the prevalence of bidirectional connections, it is directly related to the over-representation of complex triadic motifs identified in bouton-net. However, when the spectral radius exceeded 1, which generally drives the network toward the chaotic regime, the performance degradation of the two networks diverged markedly (**Fig. 3B, bottom left;** the Mann-Whitney U test, two-sided, p = 2.61e-43). Compared with the ER random network, bouton-net exhibited a substantially slower decline in memory capacity as connection weights were amplified by spectral radius, indicating reduced sensitivity to global weight scaling and enhanced dynamical stability.

This stability advantage was consistently observed in NARMA tasks involving stronger nonlinear dependencies. Although bouton-net again showed significantly lower peak performance than the random ER network across NARMA-5 (**Supplementary Fig. 2, left**; the Mann-Whitney U test, two-sided, p = 2.55e-09), NARMA-10 (**Fig. 3B, top middle**; the Mann-Whitney U test, two-sided, p = 7.51e-17), and NARMA-20 tasks (**Fig. 3B, top right**; the Mann-Whitney U test, two-sided, p = 3.97e-17), its performance remained markedly more stable in the chaotic regime (spectral radius > 1) for NARMA-5 (**Supplementary Fig. 2, right**; the Mann-Whitney U test, two-sided, p = 7.14e-205), NARMA-10 (**Fig. 3B, bottom middle**; the Mann-Whitney U test, two-sided, p = 1.36e-184), NARMA-20 (**Fig. 3B, bottom right**; the Mann-Whitney U test, two-sided, p = 3.07e-141) tasks, where the ER random network deteriorated rapidly.

Together, these results reveal a trade-off between performance and stability in biological network topology: bouton-net does not maximize task performance under optimal conditions but instead exhibits superior performance stability. This property appears to be intrinsic to its connectivity structure, particularly the elevated bidirectional connectivity, rather than arising from specific connection weights configurations.

### Complex motifs enhance network stability in bouton-net

To examine the relationship between motif occurrence and network stability, we systematically perturbed the bouton-net topology by selectively perturbing bidirectional connections (**Methods**).

Using triad census analysis, we quantified changes in motif occurrence by comparing motif counts of bouton-net against those of size-matched ER random networks. As the perturbation proportion of bidirectional connections increased, the relative occurrence of motifs M4, M5, and M8–M13 decreased monotonically (**Fig. 3C, top**). Notably, these motifs all contain bidirectional edges, confirming that the perturbation strategy directly altered the core topological features of bouton-net.

We then evaluated how these topological changes affected functional stability using the NARMA-10 task under chaotic regime (**Fig. 3C, bottom**). Within the spectral radius range of 1.0-1.4, original bouton-net consistently achieved the relatively higher task performance among all tested networks. As the perturbation proportion increased, network variants with perturbation exhibited slight linear improvements in memory performance when spectral radius was close to 1. This behavior may reflect the intrinsic sparsity of bouton-net, suggesting that its native topology does not necessarily correspond to the globally optimal stability configuration. Additionally, it also reflected that the introduction of bidirectional connections might influence the best memory task performance. However, as the spectral radius increased, network variants with higher perturbation proportion displayed more pronounced performance degradation, which was characterized by declining mean performance and increasing variance, indicating substantially reduced network stability.

These results demonstrate a strong association between network stability and the occurrence ratio of over-representation motifs. Since these motifs predominantly contain bidirectional connections, our findings suggest that the trade-off introduced by reciprocity may manifest not in maximizing computational performance, but rather in enhancing stability against dynamical perturbations. Importantly, this stability advantage emerges solely from network topology, independent of realistic connection weights, highlighting that the distinctive structural organization of biological neural networks carries intrinsic functional significance.

## Discussion

In this study, we investigated how single-neuron connectivity patterns constrain large-scale organization and dynamics in a whole-brain mouse connectome. By integrating motif analysis, modular organization, and reservoir computing, we show that the biological connectome exhibits a conserved motif-level architecture that is tightly coupled to modular directionality and dynamical stability. Rather than maximizing computational performance, the single-neuron connectome appears to prioritize stable dynamics, suggesting a fundamental trade-off embedded in its wiring principles.

First, bidirectionally connected triadic motifs are not only enriched at the whole-network level, but are consistently preserved across distinct modules composed of different brain regions. Previous connectomic studies based on electron microscopy reconstructions have reported non-random motif enrichment and reciprocal connectivity in local circuits, including insect brains and restricted mammalian cortical volumes. However, these datasets are spatially limited and do not address whether such motif organization generalizes across the entire brain. Our results extend these observations by demonstrating that motif-level complexity is a global and conserved feature of a brain-wide, single-neuron level connectome. This suggests that motif enrichment reflects fundamental constraints on biological wiring rather than region-specific circuit specializations.

Second, beyond motif abundance, we show that modular organization in the connectome is accompanied by distinct directional preferences in inter-modular connectivity. Different modules preferentially act as sources or targets of inter-modular projections, indicating asymmetric roles in large-scale information flow. Importantly, these inter-modular connections are not uniformly distributed across neurons. Instead, they are disproportionately mediated by high-degree hub neurons This observation aligns with prior work on hub nodes and rich-club organization at regional scales, and extends these concepts to the level of individual neurons in a whole-brain network. Our analysis further reveals that inter-modular projections contribute disproportionately to diverse and complex motif classes. This indicates that motifs, hubs, and modules are not independent organizational features, but are tightly intertwined. In this view, high-degree neurons serve not only as topological hubs, but also as carriers of higher-order connectivity patterns that support structured communication between modules. Such an organization may facilitate flexible and diverse routing of information while maintaining overall network coherence.

Third, we linked these structural topological features to network dynamics using a reservoir computing framework. While the biological connectome does not achieve maximal performance on memory capacity or nonlinear computation tasks under optimal conditions, it exhibits significant enhanced stability compared with random networks. In particular, performance in the chaotic regime degrades more slowly in bouton-net, indicating reduced sensitivity to the perturbation of global weight scaling. By systematically perturbing the numbers of bidirectional connections, we demonstrate that the occurrence of complex motifs, which contains bidirectional edges, is related to dynamical stability. Disrupting bidirectional connectivity reduces motif occurrence, establishing a causal connection between higher-order structural organization and emergent dynamics.

Together, our results support the view that biological neural networks are not optimized for peak computational performance, but for reliable and stable operation across a wide range of dynamical regimes. Such stability is likely essential for real brains, which must function under variable conditions, noise, and ongoing plasticity. Motif-level complexity and non-random inter-modular communication may therefore represent structural solutions evolved to balance recurrence, integration, and stability.

Several limitations of this work should be acknowledged. First, bouton-net connectivity is inferred using Peters’ rule and does not incorporate synaptic weights, cell types, or inhibitory/excitatory distinctions, which may further shape motif occurrence and dynamics. Second, reservoir computing provides a simplified and abstract proxy for neural dynamics and does not capture learning or plasticity mechanisms. Finally, while our perturbation analyses support a causal role for bidirectional motifs, the fully establishing relationships between structure and function will require future studies integrating biophysical modeling and experimental perturbations in vivo.

Despite these limitations, this study demonstrates that whole-brain, single-neuron connectomes encode conserved higher-order structure with direct functional consequences. By bridging connectomic topology and dynamical behavior, our work provides a framework for understanding how micro-scale wiring principles constrain macro-scale brain dynamics, and highlights stability as a key organizing principle of biological neural networks.

## Methods

### Random network generation and hyperparameter settings

To compare bouton-net with random networks, we selected three related and classic types of directed random networks: Erdős–Rényi (ER) network, Scale-Free (SF) network, and Stochastic Block Model (SBM) network. For each network type, 100 independent random networks were generated with random seeds ranging from 0 to 100.

For ER random networks, we used the Erdos_Renyi function (with the parameters, n=the number of neurons in bouton-net, m=the total number of edges in bouton-net, directed=True, and loops=False) from the igraph Python package (version 0.11.9), thereby ensuring that ER networks matched bouton-net in overall network size and edge count. For SF random networks, we applied the Barabasi function (with the parameters, n=the number of neurons in bouton-net, m=3, directed=True) from the igraph Python package (version 0.11.9). These parameters were chosen to be consistent with previous connectome studies [25], ensuring that the generated SF networks exhibited degree distributions correlated with that of bouton-net. For SBM random networks, we generated directed networks with 12 blocks to approximate the modular organization of bouton-net. The total number of nodes n was set equal to bouton-net, and nodes were evenly assigned to blocks, with any remainder allocated to the last block. The expected average degree of the network was matched to bouton-net, and connection probabilities were defined using an assortative stochastic block structure. Specifically, parameter α = 0.8 was used to control the fraction of edges expected to occur within blocks, yielding an inter-to intra-modular edge proportion comparable to that observed in bouton-net (4137 total edges, including 885 inter-modular edges). The resulting intra-block connection probability was higher than the inter-block probability, implemented by assigning larger probabilities to diagonal entries of the block preference matrix. The SBM networks were generated using the SBM function from igraph (version 0.11.9) with directed=True.

To ensure comparability with bouton-net, all random networks were required to be weakly connected. Connectivity was assessed using the is_connected function (with the parameters, mode=“weak”), and network generation was retried up to a maximum of 200 attempts if necessary.

### Calculation of motif occurrence ratio

We analyzed triadic motifs using the 13 directed three-node motifs: ‘021D’, ‘021U’, ‘021C’, ‘111D’, ‘111U’, ‘030T’, ‘030C’, ‘201’, ‘120D’, ‘120U’, ‘120C’, ‘210’, and ‘300’. For bouton-net, size-matched ER random networks, and topology-perturbed variants of bouton-net, motif occurrence was quantified using the get_subisomorphisms_lad function (with the parameters, induced=True) from igraph (version 0.11.9). For each motif i, we obtained the motif counts in bouton-net 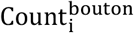 and in ER random networks 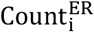. Then motif occurrence was quantified as a log-ratio relative to ER random networks, 0.1 was added to avoid division by zero:

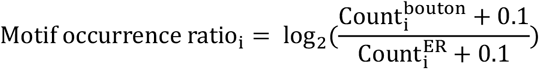

### Reservoir computing framework and parameter settings

Network dynamics were evaluated using a reservoir computing framework implemented with the Reservoir class (with the parameters, units=the number of neurons in bouton-net, Win=uniform, input_dim=1, input_connectivity=0.1, lr=1.0) from the reservoirpy Python package (version 0.4.1). For bouton-net, random networks, and topology-perturbed bouton-net variants, connection weight matrices (W) were assigned by sampling from a uniform distribution between −1 and 1, constrained by the underlying network topology. The readout layer was trained using ridge regression, implemented via the Ridge function from reservoirpy.node (version 0.4.1) with a regularization parameter of 1e-4.

### Warm-up estimation and echo-state stabilization

Before training, an adaptive warm-up period was applied to remove transient dynamics and ensure that reservoir states were independent of initial conditions. The warm-up length was determined using the test of the echo-state property (ESP), based on the convergence of reservoir trajectories initialized from different initial states under identical input sequences (the length of input sequences was 100 timesteps). For each network, the reservoir was driven by the same input sequence from two distinct initial conditions, and the Euclidean distance between the resulting state trajectories was computed at each time step and normalized by the initial distance. A reservoir was considered to have effectively forgotten its initial state when this normalized distance fell below a default threshold (ε=1e-2). The convergence time was estimated by fitting an exponential decay model to the logarithm of the trajectory distance, yielding a characteristic time constant τ. The warm-up length was then defined as:

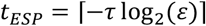

This procedure was repeated across multiple trials. If the ESP detection failed to identify convergence within the input duration in more than 10% of trials, a conservative fixed warm-up period of 100 timesteps was used. Otherwise, the warm-up length was set to the average estimated convergence time across successful trials.

### Memory capacity and NARMA tasks design

For the Memory capacity task, input sequences were generated from a normal distribution. Memory capacity was quantified as the sum of the squared Pearson correlation coefficients between the output and the input with different delays, evaluated over delays up to a maximum of *T* = 30 timesteps. For nonlinear computation tasks, we evaluated NARMA-n tasks with n=5, 10, and 20. Target sequences were generated using the narma function from reservoirpy.datasets (version 0.4.1). Network performance was quantified using the coefficient of determination (r-squared score).

### Perturbation of bidirectional connections

To assess the functional role of bidirectional connectivity, we systematically perturbed bouton-net by selectively disrupting bidirectional edges. We first identified all neuron pairs connected by bidirectional edges. For a given perturbation proportion, a subset of these pairs was randomly selected, and one directed edge from each selected pair was removed randomly. To preserve the total number of edges, a new directed edge was randomly added between two previously unconnected nodes. This procedure altered bidirectional connectivity while maintaining overall edge density.

### Outlier handling for task performance

To reduce the influence of extreme values, task performance results were filtered using a robust outlier detection procedure. For given range of spectral radius, the median and median absolute deviation were computed. Modified z-scores were calculated, and data points with absolute modified z-scores greater than 3.5 were excluded. Filtered results were then aggregated for subsequent statistical analyses and visualization.

## Acknowledgment

We thank Dr. Linus Manubens-Gil and Penghao Qian for comments on the figures and reviews on the manuscript. This work was supported by several grants awarded to H.P., who is a New Cornerstone Investigator and a Shanghai Academy of Natural Sciences (SANS) Senior Investigator. We also thank members in Peng Lab for discussion of this work.

## Author Contribution

H.P. managed this study. Y.L. developed the method and proposed the theoretical framework, and conducted the experiments. Y.W. contributed ideas of simulation part. Y.L. wrote the paper under the supervision of H.P., with input from all coauthors.

## Data and Materials Availability

All code, prompt designs, and datasets used in this study are openly accessible at the Zenodo repository (https://zenodo.org/doi/10.5281/zenodo.18627213) under the MIT License (https://opensource.org/licenses/MIT).

## Supplementary Figures

**Supplementary Figure 1.**
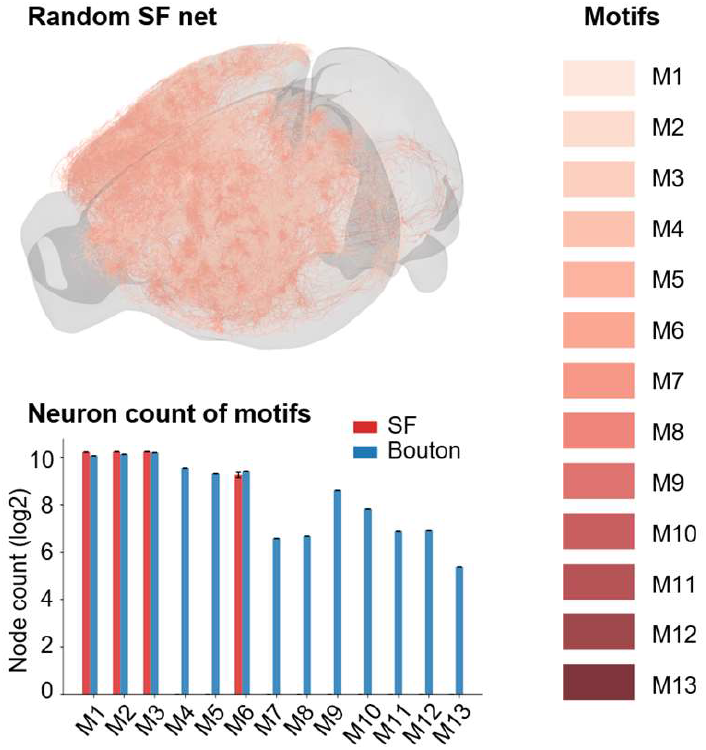
Motifs’ spatial distribution of neurons (Top) and the number of neurons which participant in motifs (Bottom) between bouton-net, Scale-Free (SF) random networks. Color bar, the level of motifs in triad census. The motif identity of each neuron is defined as its involvement of the highest motif. The error bars represent the mean neuron numbers±standard deviation.

**Supplementary Figure 2.**
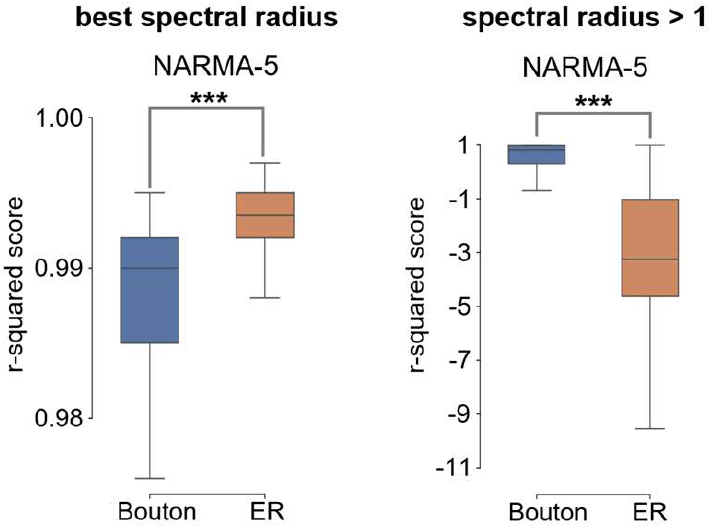
Performance comparison of NARMA-5 task between bouton net and ER random net with increasing spectral radius. Left: distribution comparison of performance at the spectral radius with best performance. Right: distribution comparison of performance with spectral radius above 1. Statistical significance was assessed using two-sided Mann–Whitney U tests, *** represents p < 0.001. Colors indicate the types of networks.

